# Microbial residents of the Atlantis Massif’s shallow serpentinite subsurface

**DOI:** 10.1101/870956

**Authors:** Shahrzad Motamedi, Beth N. Orcutt, Gretchen L. Früh-Green, Katrina I. Twing, H. Lizethe Pendleton, William J. Brazelton

**Affiliations:** School of Biological Sciences, University of Utah, Salt Lake City, Utah, USA; Bigelow Laboratory for Ocean Sciences, East Boothbay, Maine, USA; Institue of Geochemistry and Petrology, ETH Zürich, Zürich, Switzerland

## Abstract

The Atlantis Massif rises 4,000 m above the seafloor near the Mid-Atlantic Ridge and consists of rocks uplifted from Earth’s upper mantle. Exposure of the mantle rocks to seawater leads to their alteration into serpentinites. These aqueous geochemical reactions, collectively known as the process of serpentinization, are exothermic and are associated with the release of hydrogen gas (H_2_), methane (CH_4_), and small organic molecules. The biological consequences of this flux of energy and organic compounds from the Atlantis Massif were explored by International Ocean Discovery Program (IODP) Expedition 357, which used seabed drills to collect continuous sequences of shallow (<16 meters below seafloor) marine serpentinites and mafic assemblages. Here, we report the first census of microbial diversity in samples of the drill cores, as measured by environmental 16S rRNA gene amplicon sequencing. The problem of contamination of subsurface samples was a primary concern during all stages of this project, starting from the initial study design, continuing to the collection of samples from the seafloor, handling the samples shipboard and in the lab, preparing the samples for DNA extraction, and analyzing the DNA sequence data. To distinguish endemic microbial taxa of serpentinite subsurface rocks from seawater residents and other potential contaminants, the distributions of individual 16S rRNA gene sequences among all samples were evaluated, taking into consideration both presence/absence as well as relative abundances. Our results highlight a few candidate residents of the shallow serpentinite subsurface, including uncultured representatives of the Thermoplasmata, Acidobacteria, Acidimicrobiia, and Chloroflexi.

**Importance:** International Ocean Discovery Program Expedition 357: “Serpentinization and Life” utilized seabed drills for the first time to collect rocks from the oceanic crust. The recovered rock cores represent the shallow serpentinite subsurface of the Atlantis Massif, where reactions between uplifted mantle rocks and water, collectively known as serpentinization, produce environmental conditions that can stimulate biological activity and are thought to be analogous to environments that were prevalent on the early Earth and perhaps other planets. The methodology and results of this project have implications for life detection experiments, including sample return missions, and provide the first window into the diversity of microbial communities inhabiting subseafloor serpentinites.

## Introduction

Subsurface environments may host the majority of microbial life on Earth (1, 2), but current estimates of subsurface biomass lack certainty due to a paucity of observations. Recent studies, including major ocean drilling expeditions, have greatly improved the quantification and characterization of microbial populations in deep marine sediments (3–6). Seafloor crustal rocks are also expected to be an important microbial habitat, in part because a significant fraction of the oceanic crust supports hydrothermal circulation of seawater that can bring nutrients and energy into subseafloor ecosystems (7–9). The mineral composition and geological history of the host rocks determine the local environmental conditions experienced by subseafloor microbes (10–13). Ultramafic habitats for life may have unique attributes, as these rock types are unstable in the presence of seawater and undergo a series of serpentinization reactions that can support chemolithotrophy (14–16). A few microbial diversity surveys of mafic basaltic and gabbroic seafloor rocks have been conducted (17–19), but no studies have characterized the microbial diversity of ultramafic rocks.

International Ocean Discovery Program (IODP) Expedition 357 addressed this knowledge gap by targeting the Atlantis Massif, a submarine mountain located in the north Atlantic Ocean on the western flank of the Mid-Atlantic Ridge (20, 21). The massif is approximately 16 km across and rises 4,267 m from the seafloor. It is composed of variable amounts of ultramafic rocks uplifted from the upper mantle along a major fault and altered into serpentinites through the geochemical process of serpentinization (22, 23). Oxidation of iron minerals in the serpentinizing rocks results in the release of hydrogen gas and hydroxyl ions, contributing to the formation of extremely reducing, high pH fluids (24, 25). The high hydrogen concentrations that develop during serpentinization are conducive to the formation of methane and other organic compounds from the reduction of inorganic carbon (26–28). Circulation of organic-rich fluids through the serpentinites of the Atlantis Massif may support an active subseafloor ecosystem (29).

Most of our current knowledge of the biological implications of marine serpentinization comes from the exploration of the Lost City hydrothermal field, a collection of carbonate chimneys near the summit of the Atlantis Massif where high pH fluids exit the seafloor (22, 25, 30, 31). The chimneys are covered in mucilaginous biofilms formed by bacteria and archaea that are fueled by the high concentrations of hydrogen, formate, and methane in the venting fluids (32–35). The high density of the microbial biofilm communities is likely enabled by the mixing of warm, anoxic hydrothermal fluids with oxygenated ambient seawater, enabling a wide range of metabolic strategies. In contrast, the much more voluminous, rocky subsurface habitats underlying the Lost City chimneys and throughout the Atlantis Massif probably experience much more diffuse hydrothermal circulation and perhaps limited exposure to ambient seawater. Our knowledge of this subseafloor environment, however, is extremely limited (19).

The work reported here focused on two fundamental questions regarding serpentinization and life: How can we distinguish endemic microbial communities of low biomass subseafloor rocks from other environments? Which microbes inhabit marine serpentinites? To address these questions, we conducted a cultivation-independent census of microbial diversity in the shallow (<16 mbsf) rocks collected during IODP Expedition 357 (21) and compared these to the communities found in seawater and potential sources of contamination.

Laboratory contamination of DNA sequencing datasets is a well-known problem (36–39), especially for low biomass subsurface samples (19, 40–42), that must be addressed by a combination of experimental and computational methods. No matter how cleanly DNA is prepared, contaminant sequences can appear for a myriad of reasons (37)(38). Sheik *et al.* (39) recently reviewed practices for identifying contaminants in DNA sequence datasets, highlighting the importance of collecting and sequencing control samples representing potential sources of environmental and laboratory contamination. These studies have highlighted the need to complement careful laboratory protocols with computational methods that can distinguish true residents from multiple potential environmental and laboratory contaminants (43–45).

Here we report our methodological and computational strategies for minimizing and identifying contaminants in DNA sequencing datasets from extremely low-biomass seafloor serpentinites collected during IODP Expedition 357. Sequencing DNA from serpentinites required the development of a novel DNA extraction and purification protocol that recovers highly pure DNA without the use of any commercial kits or phenol. We then identified environmental and laboratory contaminants in the resulting DNA sequencing data with a computational workflow that considers the relative abundances of individual sequences among potential contamination sources. Finally, we describe the archaeal and bacterial taxa that are potential residents of seafloor serpentinites.

## Results

### DNA Yields from Rocks

DNA was extracted from at least one rock core sample collected from all sites (**Figure 1, Text S1**) except M0073 (**Table 1**, **Data set S1**). DNA yields from lysates (post-lysis, pre-purification) ranged from ∼100 to almost 500 ng of DNA per gram of rock sample. DNA yields from serpentinite and non-serpentinite rocks (including metagabbro, carbonate sand, basalt breccia and metadolerite) were similar (**Table 1 and Data set S2**). These values may be inflated by the many impurities in the lysates, which had not been purified and were visibly cloudy. No trends in DNA yield with respect to mineralogy or locations of the boreholes were apparent. During initial testing of the protocol, DNA quantification was conducted at each step of the extraction to verify that DNA levels never increased, which would indicate laboratory contamination. For the precious serpentinite samples, however, DNA was quantified only after lysis, washing, and the final purification. DNA was below the detection limit for all final, purified preparations, indicating a substantial loss of DNA from the lysates but also minimal laboratory contamination.

**Figure 1.**
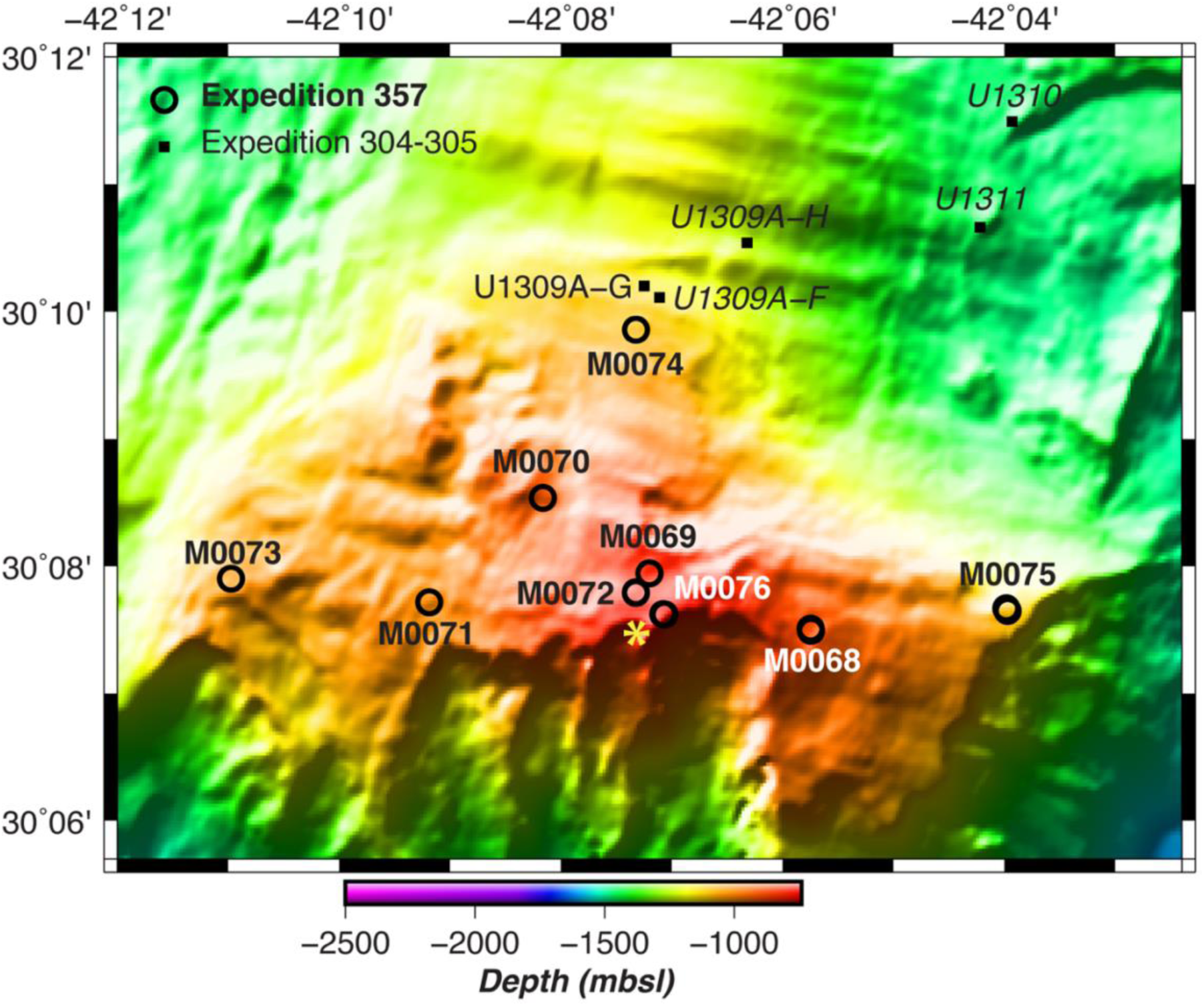
Multibeam bathymetry of the Atlantis Massif, with water depth in meters below sea level (mbsl) in color scale per legend. Drilled sites are shown as black circles. The yellow star is the location of the Lost City hydrothermal field. The distance between the most western site (M0073) and the most eastern site (M0075) is approximately 13 km (Adapted from Früh-Green, *et. al.* (20)).

**Table 1.** DNA yield from each IODP Expedition 357 serpentinite sample after lysis, washing, and purification steps. The concentration of the final, purified DNA was below the stated detection limit for each sample. Numbers of 16S rRNA gene amplicon sequences are reported before and after quality filtering.

| Sample ID | IODP ID | Extracted rock (g) | Lysate DNA (ng per g rock) | Post-washing DNA (ng per g of rock) | Purified DNA (ng per g rock) | Total sequences | Sequences after quality filtering |
| --- | --- | --- | --- | --- | --- | --- | --- |
| 0AMRd005A | 357-68B-8R-1,29-34cm | 30 | 325 | 4.07 | < 0.46 | 2985 | 1318 |
| 0AMRd014A | 357-69A-7R-1,73-75cm | 40.5 | 104 | 2.51 | < 0.55 | 2928 | 1926 |
| 0AMRd030A | 357-71B-2R-2,53-58cm | 37 | 224 | 2.34 | < 0.29 | 3893 | 2957 |
| 0AMRd031A | 357-71B-2R-1,64-66cm | 37.5 | 215 | 2.32 | < 0.49 | 2780 | 2210 |
| 0AMRd033A | 357-71C-2R-1,78-84cm | 39 | 371 | 4.99 | < 0.2 | 3920 | 3291 |
| 0AMRd034B | 357-71C-2R-1,84-90cm | 39 | 213 | 2.97 | < 0.19 | 2901 | 2294 |
| 0AMRd036A | 357-71C-5R-CC,5-10cm | 38 | 248 | 2.03 | < 0.21 | 4281 | 3442 |
| 0AMRd045A | 357-72B-3R-1,34-36cm | 38 | 473 | 15.2 | < 0.2 | 10787 | 7225 |
| 0AMRd057A | 357-75A-1R-CC,0-4cm | 26 | 136 | 2 | < 0.16 | 153,054 | 113,994 |
| 0AMRd067A | 357-76B-5R-1,14-28cm | 37 | 299 | 4.39 | < 0.22 | 4699 | 2831 |
| 0AMRd071A | 357-76B-7R-1,105-120cm | 34.5 | 100 | 7.14 | < 0.41 | 10,095 | 7742 |
| 0AMRd072A | 357-76B-9R-1,34-41cm | 37 | 110 | 2.64 | < 0.49 | 122,361 | 82,246 |
| 0AMRd073A | 357-76B-9R-1,43-52cm | 47.4 | 149 | 4.71 | < 0.82 | 5189 | 2079 |
| 0AMRd075A | 357-76B-10R-1,91-111cm | 38 | 300 | 1.49 | < 0.52 | 12,082 | 9758 |
| 0AMRd076A | 357-76B-10R-1,91-111cm | 33 | 223 | 4.3 | < 1.24 | 9399 | 7478 |

### DNA Sequencing Results

Despite the below-detection levels of purified DNA, a total of 1,125,191 sequencing read pairs of the 16S rRNA gene were obtained from all rock samples (50 samples including sequencing duplicates). Here we focus only on the DNA sequencing results from the 15 samples characterized as serpentinites, but the DNA sequencing data from all samples are included in our SRA BioProject PRJNA575221. At least 2,780 read pairs were obtained from each sample, and >10,000 read pairs were obtained from five of the serpentinite rock samples (**Table 1**). We identified 2,063 amplicon sequence variants (ASVs) among all rock samples, 664 of which were found only in serpentinites. We obtained 27,524,256 read pairs from the water samples (76 samples of surface, shallow, and deep including sequencing duplicates) (**Data set S3**), from which we identified 15,142 ASVs. Six samples of lab air yielded 66,846 read pairs and 293 ASVs (**Data set S4**), despite below-detection levels of purified DNA.

The proportional abundances of all 17,081 unique ASVs among all rock, water, and air samples were used to calculate the community dissimilarity between each pair of samples. The results are visualized in the non-metric multidimensional scaling (NMDS) plot in **Figure 2**, which shows a clear split between water and rock samples. The bacterial communities of both serpentinite (orange points) and non-serpentinite (yellow points) rock samples are highly variable and are not consistently distinct from each other. The bacterial compositions of lab air samples (green points) overlap with those of the rock samples in this visualization.

**Figure 2:**
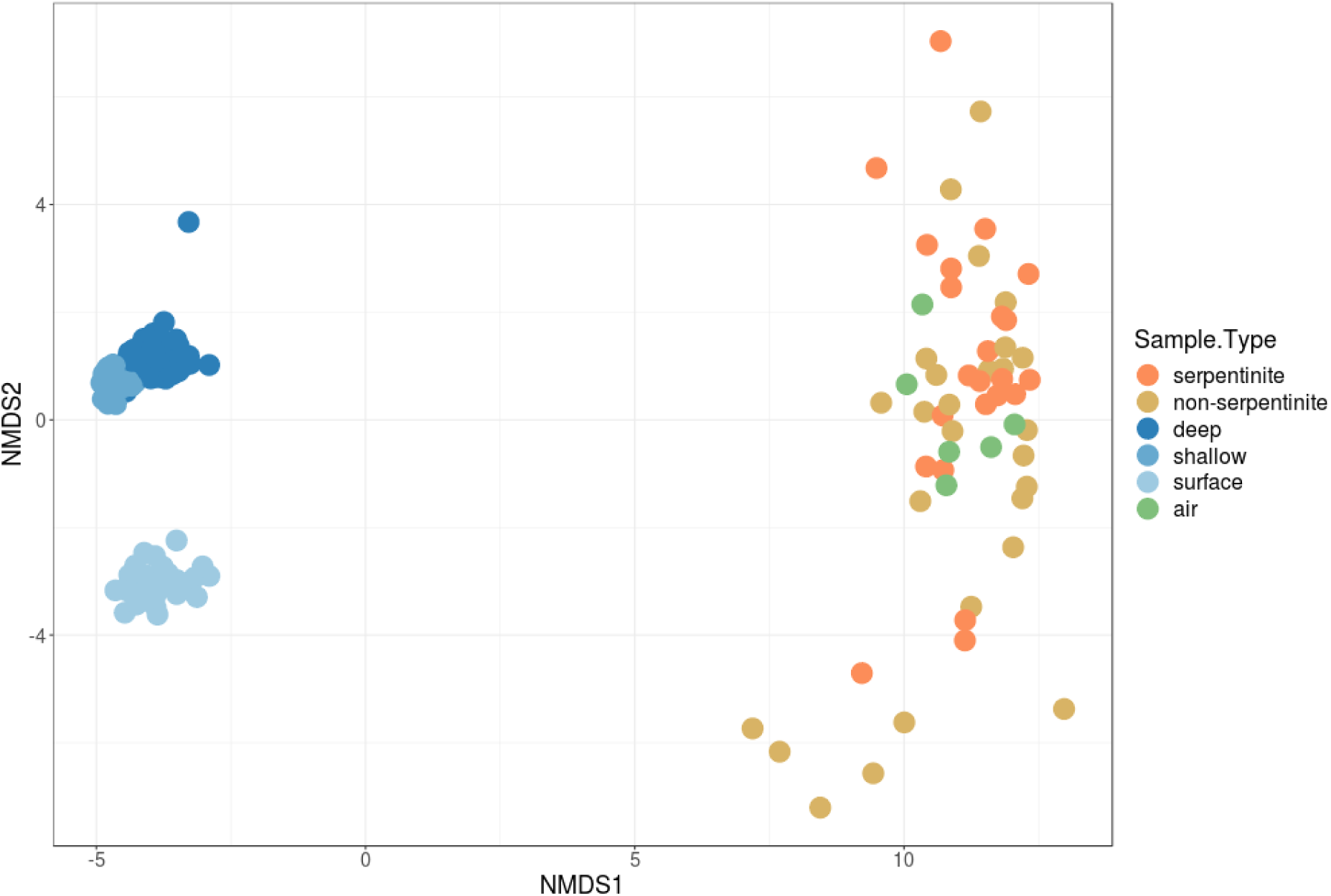
A non-metric multidimensional scaling plot shows general patterns in the 16S rRNA gene microbial community compositions of all types of samples. Each data point represents the microbial community composition of a sample of seawater (deep, shallow, or surface), rock core (serpentinite versus non-serpentinite), or laboratory air. The distances between the data points represent the Morisita-Horn dissimilarities among the microbial communities.

### Evaluation of Contamination Sources

To evaluate contamination in the serpentinites, we used SourceTracker2 (45) to estimate the proportion of each rock sample’s DNA sequences that could be attributed to contamination from water or air. These results show that very few of the DNA sequences in serpentinites can be traced back to seawater (**Figure 3**). The number of sequences attributed to air was highly variable among the serpentinites, from nearly zero to >50% of total sequences. Most serpentinite samples were dominated by sequences that could not be attributed to a single source by SourceTracker2. These ASVs are designated as having an “unknown” source and may represent true inhabitants of the serpentinites or may be derived from another source that was not sampled in this study. Based on these results, serpentinite samples with >15% of sequences attributed to lab air were excluded from further analyses (0AMRd014A, 031C, 033A, 033C, 034A, 036A, 045B, 067C, 071A, and 073A).

**Figure 3:**
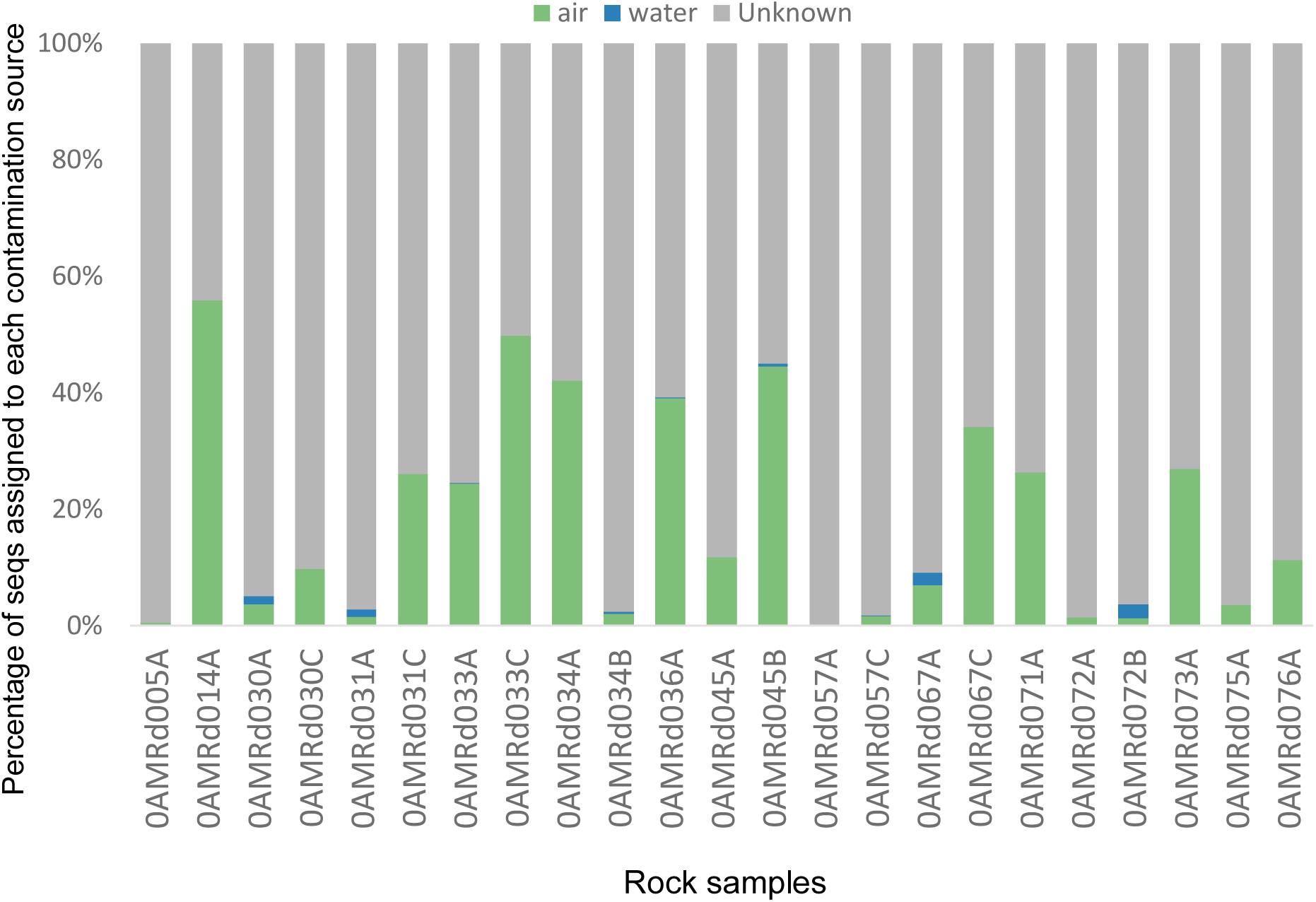
Assessment of potential 16S rRNA gene library contamination in serpentinite samples with SourceTracker2. Each bar on this plot shows the percentage of ASVs from each source of contamination into each serpentinite sample. Green = lab air. Blue = seawater samples. Gray = undetermined.

SourceTracker2 computes probabilistic estimates of how the whole-community composition of samples relate to each other, but it is not designed to identify the source of each taxon. For example, multiple ASVs in sample 0AMRd057A occurred in both air and water samples, which some approaches would consider to be evidence of contamination, even though SourceTracker2 estimated that <1% of sequences in this sample could be assigned to water or air sources (**Figure 3**). Therefore, we explored other methods for the identification of individual ASVs as contaminants.

### Identification of Contaminants from Air and Seawater

Of the 293 ASVs detected in air samples, 187 also occurred in serpentinites and were therefore designated as contaminants (**Figure 4A**). Thus, all remaining ASVs were absent in air and were only detected in water and serpentinite samples. Next, two approaches, simple overlap (SO) and differential abundance (DA), were used in parallel to distinguish the serpentinite ASVs from the water ASVs. In the SO approach (**Figure 4B**), the ASVs shared between water and serpentinite samples were removed from the dataset regardless of their abundances in water or serpentinites (i.e. only considering presence/absence). Thus, the results from the SO approach include only those ASVs that are exclusively found in serpentinite samples (664 ASVs).

**Figure 4:**
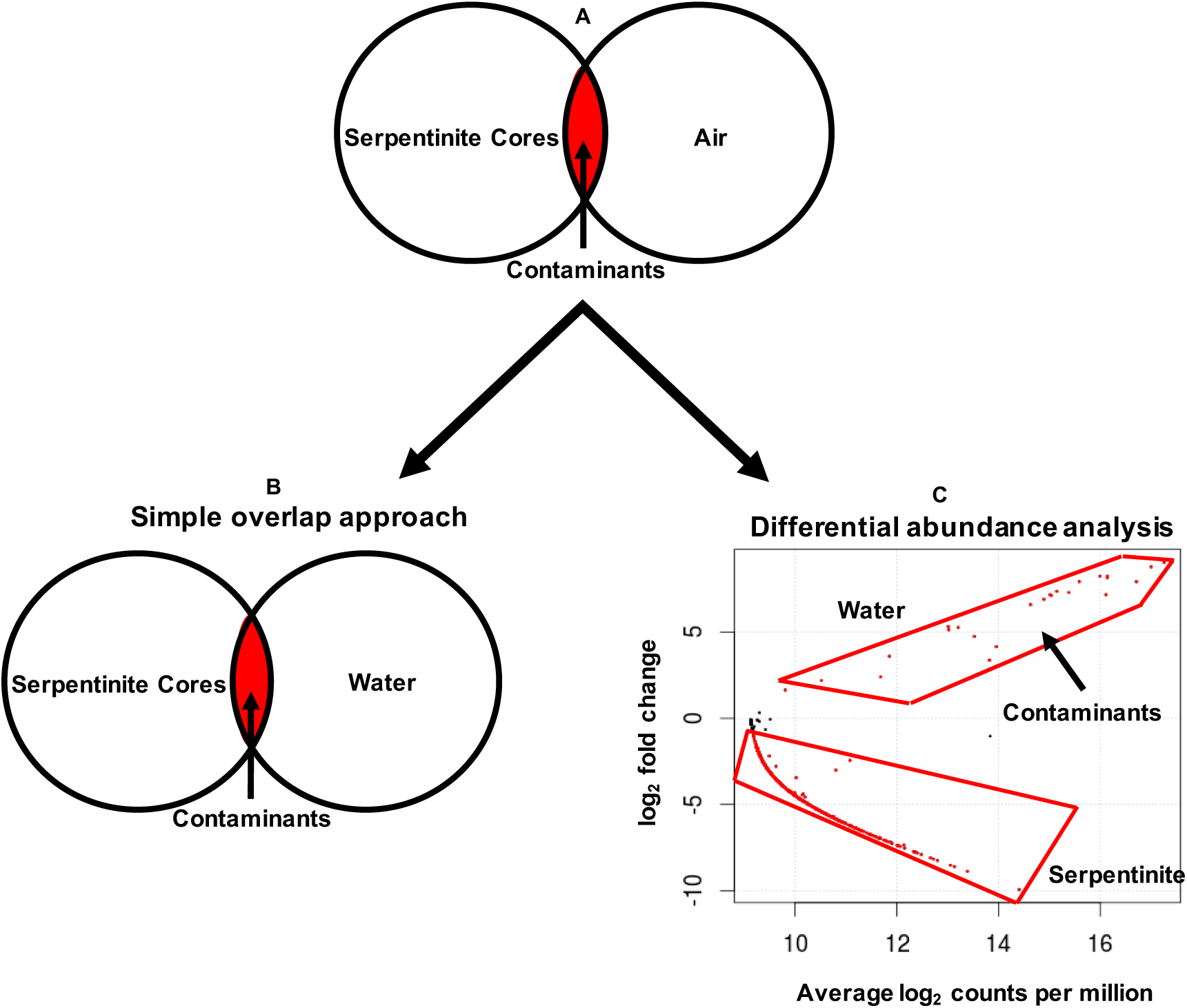
Workflow for identifying DNA sequence contaminants. A) All ASVs detected in air samples were removed from the dataset. B) The Simple Overlap (SO) approach was used to remove all ASVs detected in water samples. C) The Differential Abundance (DA) approach was used to remove only those ASVs that were significantly more abundant in water samples compared to serpentinite samples. Differentially abundant ASVs are shown as red data points in the plot. The significance threshold was a false detection rate of 0.05. Only one of the comparisons is shown here (deep water samples vs. serpentinite samples). Shallow and surface water samples were compared in separate analyses not shown here.

In the differential abundance (DA) approach (**Figure 4C**), the sequence counts (i.e. the number of merged paired reads) of each ASV shared between water and serpentinite samples were compared. ASVs that were significantly more abundant in water samples compared to serpentinites were identified and then removed from the dataset (**Table S3**). In this case, ASVs that are highly abundant in serpentinites and very rare (but still present) in water are not necessarily identified as contaminants, as they were in the SO approach. Three edgeR comparisons were performed between serpentinite samples and each depth category of water samples (serpentinites vs. surface, serpentinites vs. shallow, and serpentinites vs. deep). The ASVs that were more abundant in each group of water samples (one example in **Figure 4C**) were designated as contaminants from water and excluded from further analyses. The ASVs identified as water contaminants and detected in serpentinites are listed in **Data set S5**. The DA approach results in a final list of 684 ASVs putatively considered to be true inhabitants of serpentinites (**Data set S6**), including 20 additional ASVs compared to the SO approach. Seven of these 20 ASVs were present at >100 counts in serpentinite samples and less than a total of 53 counts in all water samples (**Data set S7)**.

### Taxonomic Composition of Serpentinites

To conservatively report a shorter list of likely serpentinite inhabitants, the following analyses and visualizations only include ASVs with at least 100 total counts among all samples. Taxonomic classifications of the putative serpentinite ASVs from the SO approach are summarized in **Figure 5**. For clarity, the taxonomic classifications of only the top 50 most abundant ASVs are shown, and more information about these 50 ASVs is provided in **Data set S7**. The taxonomic compositions of serpentinite samples are variable, consistent with the spread of serpentinites shown in the NMDS plot (**Figure 2**). No significant differences with respect to sample handling (flamed/unflamed, shaved/unshaved, washed/unwashed) could be detected. Indeed, very few similarities between any two samples were apparent.

**Figure 5:**
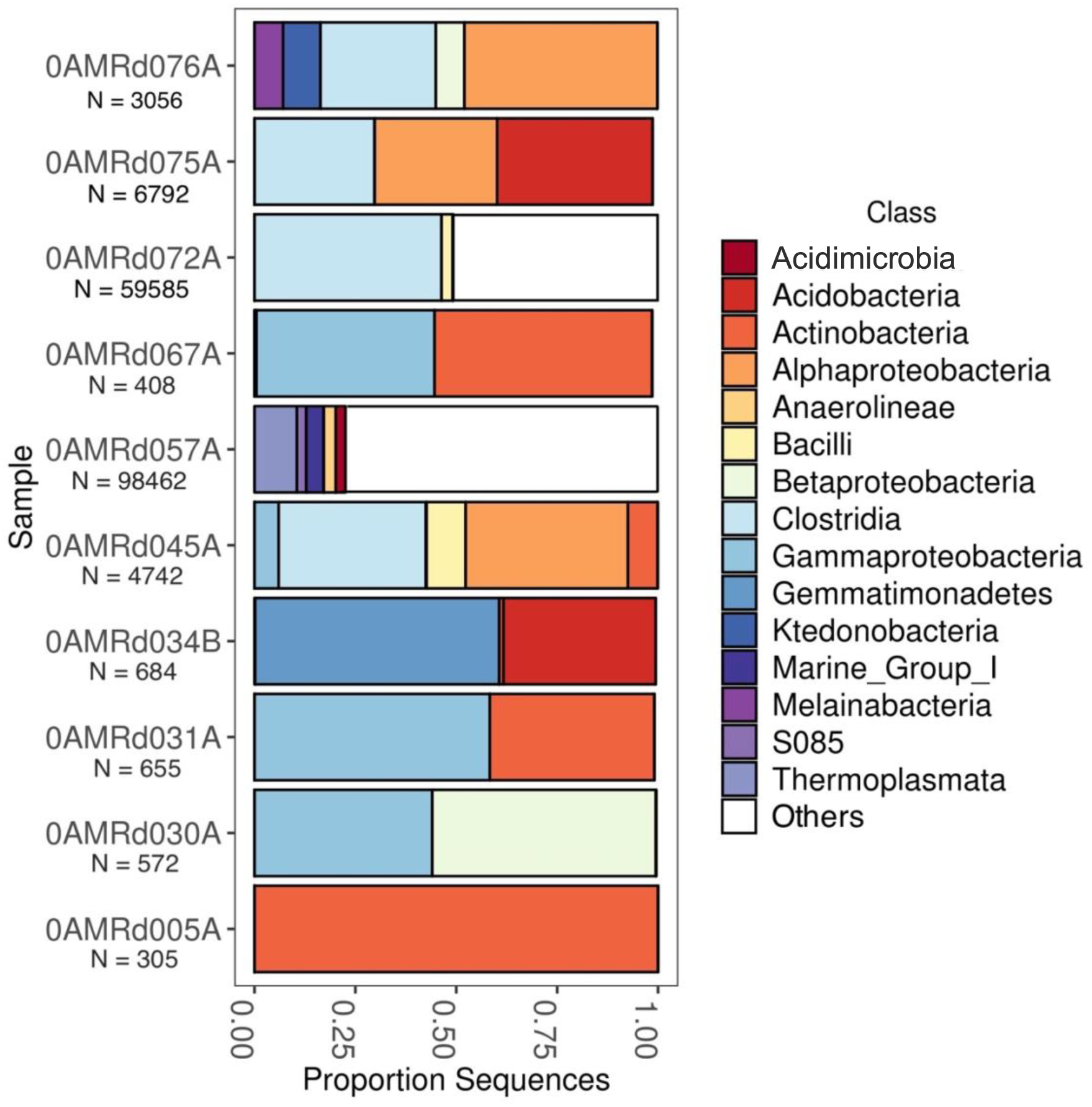
Taxonomic summary of ASVs detected in serpentinite samples after contaminant removal. For clarity, only the top 50 ASVs among all samples (Supplementary Table S8) are shown here. N is the total ASV counts for each sample after excluding rare ASVs. Samples 0AMRd076A, 075A, 072A, and 067A were collected from hole M0076B. Samples 0AMRd031A and 030A were collected from hole M0071B. No other samples in this figure were collected from the same hole.

Sample 0AMRd057A (**Figure 6**) provided the deepest sequencing dataset (98,462 merged paired reads among those ASVs with at least 100 counts). Most of the taxonomic diversity in this sample was represented by rare ASVs (i.e. those with lower abundance than the top 50 shown in **Figure 5**). The most abundant ASV was classified as family Thermoplasmatales (phylum Euryarchaeota) and has 99% identity to a clone (NCBI accession number: AB825685, **Data set S7**) from a deep-sea sediment core in the Okinawa Trough (46). Two of the top ASVs in this sample were classified as two different classes of Chloroflexi (S085 and Anaerolineae), each of which was unique to sample 0AMRd057A and was most similar to clones from marine sediments (AM998115; HQ721355). Another top ASV unique to this sample was classified as Acidimicrobia clade OM1 (phylum Actinobacteria), which was nearly identical to a clone from a marine endolithic community (47). Two ASVs in 0AMRd057A were classified as Acidobacteria Subgroup 21 and were most similar to sequences from marine sediments, including a clone (KY977768) from altered rocks of the Mariana subduction zone, where serpentinization also occurs (48).

**Figure 6:**
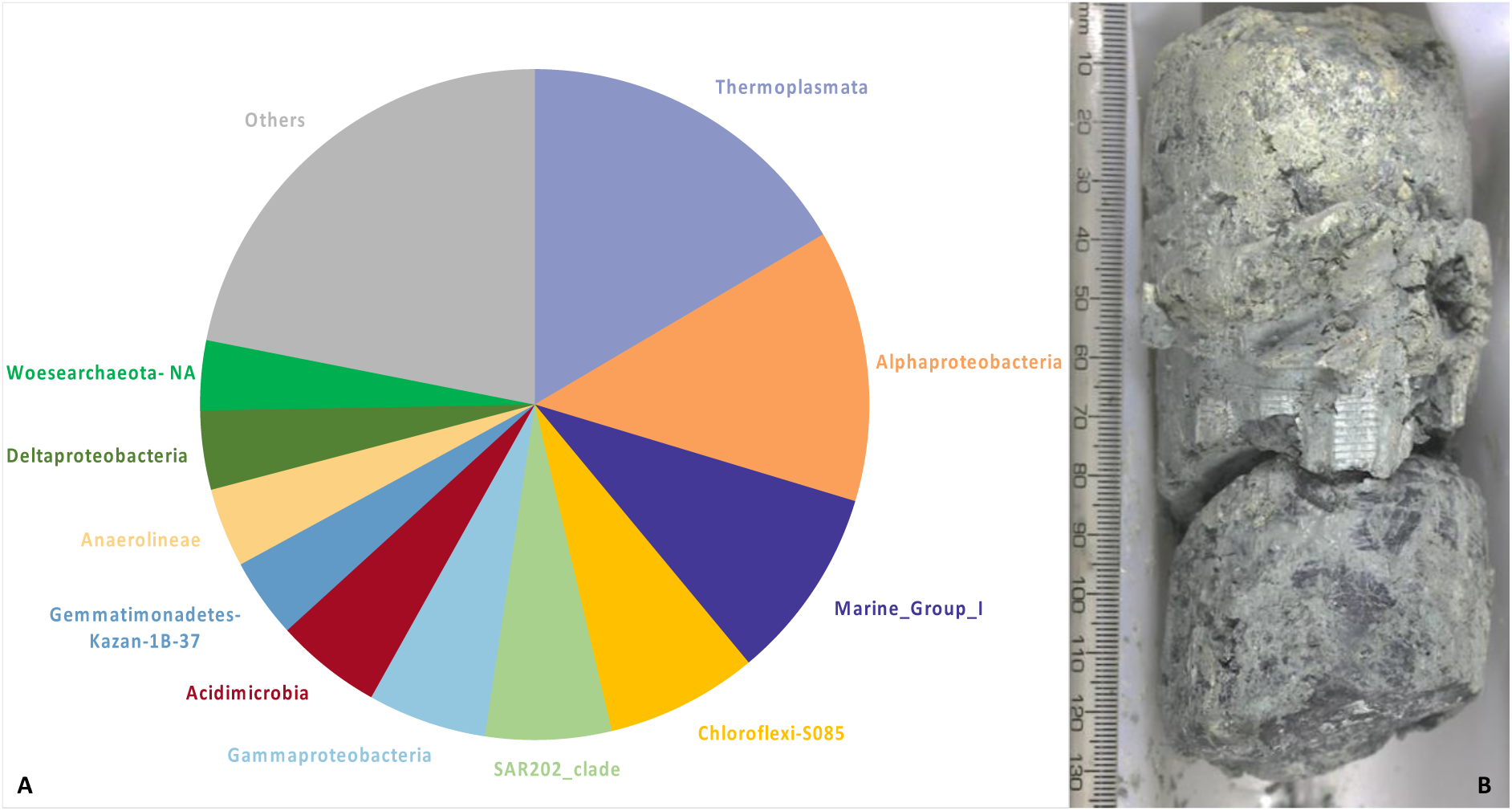
A) Taxonomic summary of serpentinite sample 0AMRd057 (357-75A-1R-CC,0-4cm). The relative abundance of each taxonomic group is its proportion of 103,229 total sequence counts among the final 331 ASVs identified by the differential abundance (DA) approach. B) This sample was recovered from the core catcher section of the core from borehole M0075A. Scale is in millimeters (71).

The deepest sequencing datasets of the remaining samples were obtained from 0AMRd045A, 0AMRd072A, 0AMRd075A, and 0AMRd076A, which were collected from the sites closest to the Lost City chimneys (boreholes M0072B and M0076B; **Figure 1**). Each of these samples were dominated by Clostridia and Alphaproteobacteria (**Figure 5**), and most of these ASVs match sequences associated with animal digestive tracts or soil. An exception is a *Rhodobacter* ASV unique to 0AMRd045A that matches several sequences from lakes (e.g. MF994002). An abundant Sphingomonadaceae ASV in 0AMRd075A matches a clone from groundwater near the Hanford Site, WA (KT431076). A *Dechloromonas* ASV unique to 0AMRd076A (Site M0076) is identical to a clone from an ocean drilling expedition to the South Chamorro Seamount (LC279322), which is also a site of subseafloor serpentinization (49). However, the genus *Dechloromonas* has also been detected on human skin (50).

Sample 0AMRd030A is dominated by two ASVs classified as genera *Acidovorax* (class Betaproteobacteria) and *Cobetia* (class Gammaproteobacteria), respectively. The *Acidovorax* ASV matches sequences from soil (e.g. CP021359) and an anoxic lake (KY515689)(51). *Cobetia marina* (previously *Halomonas marina*) is a ubiquitous marine bacterium (52) and also a model organism for the study of biofouling due to its ability to grow as a biofilm on a variety of surfaces (53).

Other abundant ASVs shown in **Figure 5** and listed in **Data set S7** include those classified as Acidobacteria, Actinobacteria, Betaproteobacteria, Gemmatimonadetes, and Ktedonobacteria, and these sequences were generally most similar to environmental sequences from soils or freshwater. One possible exception is an ASV that was abundant in sample 0AMRd031A and was classified as genus *Modestobacter*, a soil bacterium, but it was also identical to another species in the same family, *Klenkia marina*, which was isolated from marine sediment (NR_156810).

## Discussion

### Serpentinite samples have minimal environmental contamination

Sequencing of environmental DNA from seafloor rocks is challenging due to their very low biomass and their abundance of minerals (e.g. phyllosilicates) that inhibit enzymatic reactions and bind DNA (54–56). The extremely low biomass of seafloor rocks also makes them susceptible to environmental and laboratory contamination. Many precautions were taken during handling of the rock core samples to minimize, eliminate, and detect contamination, and our results agree with previous reports that these efforts were generally successful (20, 57, 58).

To assess environmental contamination of the cores, perfluoromethylcyclohexane (PFC) was injected into the seabed drilling fluids during coring. Orcutt *et al*. (57) reported that the interior zones of rocks whose outer surfaces had been shaved off with a sterile rock saw had greatly reduced levels of PFC. Only one sample in our study (0AMRd073), whose exterior had been shaved, contained detectable levels of PFC. Two additional samples (0AMRd030, 0AMRd036), which were too rubbly to be shaved with the rock saw and were instead washed with ultrapure water, contained detectable PFC. In general, however, washing rubbly samples with ultrapure water was largely effective in reducing the level of PFC, and presumably of any other surface contamination (57). The PFC tracer, however, was not designed to track any seawater contamination that might have occurred after drilling or any contamination that might have resulted from handling the rock cores shipboard or in the laboratory.

We assessed environmental contamination of the serpentinites by sequencing DNA from samples of seawater collected above the boreholes. The contribution of seawater DNA sequences into the serpentinites was minimal (**Figure 3**), further demonstrating that the sampling-handling precautions on the ship and at the Kochi Core Center were successful in eliminating environmental contamination that must have occurred at some level during core recovery on the seafloor. In addition, our results agree with those of Orcutt *et al.* (57) that there was no noticeable difference in contamination levels between rocks whose outer surfaces had been subjected to flame sterilization compared to those that were not flamed.

### Serpentinite samples have minimal environmental DNA

Our yields of ∼200 ng of DNA per g of rock sample in the crude lysates (**Table 1**) were much higher than expected from the previously reported numbers of cells for these serpentinites (20), all of which were only slightly greater than the quantification limit of ∼10 cells cm^-3^. Such a cell density would predict 20 fg of DNA cm^-3^ (assuming 2 fg of DNA per cell), which is approximately 7 orders of magnitude less DNA than observed. Therefore, the total environmental DNA must have been primarily extracellular. Nearly all of the DNA was lost during purification and fell below our detection limit, so much of it was probably highly fragmented. Indeed, neither the filtration units used for washing nor the electrophoretic purification method are expected to recover DNA below 600 bp.

Our lowest detection limit of purified DNA (which was concentrated from many parallel extractions of 30-40 grams of rock per sample) was 0.05 ng per gram of rock, which is still 10,000 times greater than that expected from the cell numbers. However, amplicon sequencing of 16S rRNA genes in the environmental DNA was moderately successful, most likely thanks to the elimination of PCR inhibitors by the electrophoretic SCODA purification. Because of the exceedingly low levels of input DNA, the probability of contamination during purification, PCR, and sequencing was very high. Indeed, laboratory contamination was evident in the serpentinites, as represented by their relatively high proportions of DNA sequences from samples of our laboratory’s air (**Figure 3**).

### Identification of laboratory contaminants

We identified laboratory contaminants as amplicon sequence variants (ASVs) that occurred in samples of both air and serpentinites. This simple overlap (SO) approach is commonly used in environmental microbiology studies to remove all sequences (or taxonomic categories) that are shared between any sources of contamination and the sample of interest (39). This approach can be useful for removing ASVs from obvious sources of contamination. An important assumption of the SO approach is that the direction of contamination is from the suspected contamination source into the sample. However, this assumption is often untested. For example, if a low-DNA control sample is handled in close proximity (e.g. the same 96-well plate) to a high-DNA environmental sample, it may be more likely for “contamination” to occur from the environmental sample into the control sample. Subsequent use of the SO approach would erroneously identify an abundant member of the environmental sample as a laboratory contaminant.

We considered the possibility of “reverse contamination” from our core samples into our air samples to be unlikely for multiple reasons. First, most of the processing of the rock samples, including sawing, crushing, washing, and subsampling, occurred at the Kochi Core Center in Japan, and our samples of air were collected in our laboratory at the University of Utah. Second, our SourceTracker2 results implicated air as a major source of DNA into the serpentinites (**Figure 3**). Third, nearly all of the taxa detected in the air samples are typically associated with humans (e.g. *Pseudomonas*, *Staphylococcus*, *Acinetobacter*). Our list of laboratory air taxa (**Data set S4**) may be useful for identifying contaminants in future studies, as a complement to previously reported lists of contaminants from laboratory reagents (37–39).

### Identification of environmental contaminants

Identification of environmental contaminants from seawater into the serpentinites was more complex because of the greater possibility of mixing between seawater and rocks in both directions. The Lost City chimneys provide dramatic evidence of the flux of biological material from the rocky subsurface into ambient seawater (25), and flux from the boreholes in this study was also evident by the release of bubbles during drilling (20). Deep seawater near the Atlantis Massif cannot be assumed to be completely free of subsurface inhabitants, and samples of seawater at shallower depths or locations far removed from the Atlantis Massif would not be accurate representations of the environmental contamination that the serpentinite samples experienced during recovery on the seafloor. Therefore, the relative abundances of taxa in samples of deep seawater and serpentinites should be informative because true subsurface organisms that are mixed into ambient seawater should be immediately diluted and present in seawater only at very low abundances.

In such situations of environmental mixing, the SO approach can erroneously identify contaminants because it ignores abundance information. We identified at least one such example in our study by comparing the results of the SO approach with the differential abundance (DA) approach. An ASV classified as Acidobacteria Subgroup 21 occurred 264 times in a serpentinite sample (0AMRd057A; 0.3% of total sequence counts in that sample) from borehole M0075 and also occurred a total of 12 times among five deep seawater samples collected from above boreholes M0068, M0072, and M0073 (**Data set S6**). The ASV was identical to several sequences from marine sediments as well as one from altered rocks of the Mariana subduction zone (KY977768) and another from seafloor lavas near the Loi’hi seamount (EU491098)(17).

This was the most striking example in our study, but there were 19 additional ASVs flagged as contaminants by the SO approach but not by the DA approach. This relatively small difference (664 compared to 684 total ASVs in serpentinites) is probably a consequence of the success in minimizing and eliminating environmental contamination during handling of the rock cores, as discussed above and demonstrated by the minimal level of contamination from seawater shown in **Figure 3**. Differences between the SO and DA approaches are likely to be greater in studies where environmental contamination cannot be controlled so carefully.

### Identification of contaminants by taxonomy

After removal of DNA sequences identified as contaminants from seawater and air, the most abundant serpentinite ASVs (**Data set S6**) still contained many taxa that seemed unlikely to be true inhabitants of the subseafloor. Some are typically associated with animal digestive tracts (e.g. *Ruminiclostridium*), some are found in soil (e.g. various Acidobacteria), and others have been previously identified as reagent contaminants (e.g. *Micrococcus*)(37, 38). The sources of these likely contaminants could not be determined in this study, but the extraction and sequencing reagents are strong candidates, even though we took great care in preparation of our own reagents at every step and did not use any commercial extraction or purification kits. Also, we only sampled and sequenced DNA from the air in our own laboratory at the University of Utah and did not sample the air inside the research vessel, the Kochi Core Center, or the DNA sequencing core facility at Michigan State University.

Identification of contaminants at the level of taxonomic category is risky, however, because organisms that are capable of persisting in the extreme environments of mostly sterile laboratory supplies, may also be capable of inhabiting the extreme environments of mostly sterile subsurface rocks (59). For example, a relatively abundant serpentinite ASV that was absent in all seawater and air samples is nevertheless a suspected contaminant because it was classified as *Acidovorax*, a ubiquitous soil bacterium and plant pathogen that has also been identified as a contaminant in laboratory reagents (38). The ability of *Acidovorax* to form biofilms and perform anaerobic iron oxidation (60), however, would be advantageous in the serpentinite subsurface. Because we are suspicious of these taxa but lack direct experimental evidence of a source other than serpentinites, we include these ASVs in our final tables but flag them as suspected contaminants (**Data set S7**).

### Putative inhabitants of the serpentinite subsurface

Excluding all air and seawater ASVs and ignoring all suspicious taxa leaves us with a few putative inhabitants of subseafloor serpentinites (**Data set S7**). Many of the these ASVs were unique to a single serpentinite sample shown in **Figure 6A**. This serpentinite was recovered from the most eastern site (borehole M0075A) at a depth of ∼60 cm below the seafloor, and it had extensive talc-amphibole alteration, suggesting that it was eroded from the central dome during uplift of the massif. The most abundant taxa in this sample included Euryarchaeota, Chloroflexi, Actinobacteria, and Acidobacteria, all of which matched DNA sequences previously found in marine sediments and rocks. The most abundant ASV was classified as Thermoplasmatales (an order within the Euryarchaeota) and was nearly identical to a clone from hydrothermal sediment collected during IODP Expedition 331 to the Mid-Okinawa Trough (46). Most, but not all, of the characterized members of the Thermoplasmatales are thermophilic, and they are typically involved in sulfur cycling (61, 62). The Thermoplasmatales are generally known as acidophiles (63), which is curious since serpentinization is associated with high pH.

The most abundant taxa in serpentinites recovered from the central dome of the massif (boreholes M0069, M0072, and M0076) were more difficult to interpret. At least two probable subsurface taxa (Sphingomonadaceae and *Dechloromonas*) were detected in these serpentinites, but many of the other sequences from these samples are identical to environmental sequences from studies of animal digestive tracts or soils. Therefore, serpentinites from the central dome, which are considered to be most representative of the massif’s basement, appear to be more susceptible to undetected contamination, perhaps due to extremely low DNA yields from these samples.

Recently, Quéméneur *et al*. (58) reported enrichment cultures obtained from IODP Expedition 357 rock cores, including two serpentinite samples. One of these was a section of a serpentinite core (357-71C-5R-CC) that was adjacent to a sample included in the present study (0AMRd036A). Our parallel sample was heavily contaminated with air DNA (**Figure 3**) and did not contain any of the dominant taxa detected by Quéméneur *et al*. Many of the taxa detected in their cultivation experiments were also present in our results, but only one genus reported by Quéméneur *et al*. (*Sphingomonas*) was included in our final list of candidate serpentinite taxa (**Data set S7**). Most of the successful enrichments reported by Quéméneur *et al*. were obtained from a carbonate-hosted basalt breccia, which was not included in our study.

In 2005, IODP expedition 304/305 recovered gabbroic rocks from the central dome of the Atlantis Massif, but no serpentinites (**Figure 1**). A few phylotypes of proteobacteria were detected in the gabbroic rock cores by sequencing of environmental 16S rRNA genes (19). None of these matched the candidate serpentinite taxa reported here. However, most of the functional genes detected by Mason *et al*. (19) with the GeoChip microarray, including genes associated with metal toxicity, carbon degradation, denitrification, and carbon fixation, are consistent with the taxa detected in our serpentinite samples.

## Conclusions

This study provides the first sequences of environmental DNA from subseafloor serpentinites, enabled by the high recovery of rock cores by IODP Expedition 357 to the Atlantis Massif (20). The extremely low biomass of the serpentinites presented multiple challenges, necessitating the development of a novel DNA extraction and purification protocol. We developed strategies to increase the yield of DNA appropriate for PCR amplification, while eliminating many potential sources of laboratory contamination. We hope that our methodology, as well as our identification of laboratory contaminants derived from dust particles, will prove to be useful to other researchers studying extremely low-biomass environments. We were able to identify candidate residents of the serpentinite subseafloor by employing a series of contamination-detection procedures, including evaluations of whole-sample compositions and of individual sequences. Our results highlight the importance of a multi-faceted approach to contamination control that features sampling of potential contamination sources as a complement to microbiologically clean sample-processing protocols. Multiple statistical and bioinformatic tools were used to identify contaminant DNA sequences in a careful process that did not unnecessarily ignore useful information, such as the relative abundances of individual DNA sequences in potential contamination sources. No computational tool can ever prove that an environmental DNA sequence is not a contaminant, and future studies will investigate candidate subseafloor organisms with experimental studies.

## Materials and Methods

Detailed descriptions of samples and sample-processing protocols are available in the Supplemental Methods (**Text S1**). DNA was purified from rock samples with SCODA (synchronous coefficient of drag attenuation) technology implemented with the Aurora purification system (Boreal Genomics, Vancouver, British Columbia, Canada). 16S rRNA gene amplicon sequencing was conducted by the Michigan State University genomics core facility on all of the samples (rock, water, and lab air). All samples were sequenced twice (i.e. sequencing replicates). The V4 region of the bacterial 16S rRNA gene was amplified with dual-indexed Illumina fusion primers (515F/806R) as described elsewhere (64). 16S rRNA gene amplicon sequences were processed with cutadapt v. 1.15 (65) and DADA2 v. 1.10.1 (66). Taxonomic classification of all ASVs was performed with DADA2 using the SILVA reference alignment (SSURefv132) and taxonomy outline (67). Raw counts were converted to proportions to normalize for variations in sequencing depth among samples. The non-metric multi-dimensional scaling (NMDS) plot was generated from a table of ASV proportional abundances across all sample categories (rocks, water, lab air) using the distance, ordinate, and plot_ordination commands in the R package phyloseq v.1.26.1 (68). The major sources of contamination and the level of contamination in each sample were estimated with SourceTracker2, version 2.0.1 (45). Differential abundance was tested with the R package edgeR v. 3.24.3 (69) as recommended elsewhere (70). We used edgeR to contrast the total read counts of ASVs in serpentinite rock samples compared to three groups of water samples (surface, shallow, deep). ASVs that were absent in all serpentinites and ASVs with low variance (<1e-6) were excluded from the comparisons. The output of the three edgeR tests (serpentinite samples compared to each of the three groups of water samples) was three lists of ASVs with significant differential abundances (false discovery rate (FDR) < 0.05) in serpentinite samples or water samples. The final list of ASVs was created by deleting ASVs with greater abundances in any of the categories of water samples (as determined by the edgeR tests) from the original list of ASVs from which all air ASVs had already been removed. Finally, rare ASVs (those that did not have >=100 counts in a single sample) were excluded from the final results, merely as a conservative abundance filter for reporting a final list of ASVs expected to be present in serpentinite rocks. No comparisons of diversity were attempted after removing rare ASVs.

## Data Availability

All unprocessed DNA sequence data are publicly available at the NCBI SRA under BioProject PRJNA575221. All supplementary data and protocols are available at https://github.com/Brazelton-Lab/Atlantis-Massif-2015. All custom software and scripts are available at https://github.com/Brazelton-Lab.

## Acknowledgements

This research used data and samples provided by the International Ocean Discovery Program (IODP) Expedition 357 supported by the European Consortium for Ocean Research Drilling (ECORD) and implemented by the ECORD Science Operator (ESO). The shipboard sample-processing protocols were executed by ESO Expedition Project Managers Carol Cotterill and Sophie Green, ESO Operations Superintendent David Smith, the crews of the R.R.S. *James Cook* and the MeBo and RD2 seabed drills, and shipboard scientists Susan Lang, Marvin Lilley, Yuki Morono, Marianne Quéméneur, and Matthew Schrenk. We are indebted to Fumio Inagaki for welcoming our use of sampling equipment at the JAMSTEC Kochi Core Center (KCC), to Nan Xiao at KCC for coordinating our sample processing and operation of core sectioning equipment, to Christopher Thornton, Katherine Hickock, and Susan Lang for manually processing the rock samples at KCC, and to ESO staff Ursula Röhl and David McInroy for helping to coordinate sample transfer and tracking. Christopher Thornton, Alex Hyer, Emily Dart, and Julia McGonigle provided laboratory and computational assistance. We also thank Jackie Goordial for comments on an earlier draft. The Expedition 357 core description team helpfully provided descriptions of the core lithologies. Funding for this work was provided by the NASA Astrobiology Institute Rock-Powered Life team, the U.S. National Science Foundation-funded US Science Support Program (subawards to WJB and BNO), the Deep Carbon Observatory funded by the Alfred P. Sloan Foundation (via subaward to BNO from the Marine Biological Laboratory), and by the Swiss National Science Foundation (SNSF) project No. 200021_163187 (to GFG) and SNSF contributions to the Swiss IODP.

## Supplemental files

**Data set S1**: Names and locations of all serpentinite and non-serpentinite rock samples.

**Data set S2**: DNA yield from non-serpentinite rock samples.

**Data set S3**: Names and locations of all seawater samples.

**Data set S4**: List of ASVs detected in samples of laboratory air.

**Data set S5**: List of ASVs detected in samples of seawater.

**Data set S6**: Final list of 684 ASVs in serpentinites determined by the DA method.

**Data set S7**: Top 50 most abundant ASVs in serpentinites (according to maximum proportional abundance per sample).

**Text S1**: Supplementary descriptions of samples and sample-processing methodology.

